# Sarcoptic mange outbreak decimates South American camelid populations in San Guillermo National Park, Argentina

**DOI:** 10.1101/2021.08.18.456827

**Authors:** Hebe Ferreyra, Jaime Rudd, Janet Foley, Ralph E. T. Vanstreels, Ana M. Martín, Emiliano Donadio, Marcela M. Uhart

## Abstract

Sarcoptic mange epidemics can devastate wildlife populations. In 2014, mange was first detected in vicuña (*Vicugna vicugna*) and guanaco (*Lama guanicoe*) in San Guillermo National Park (SGNP), Argentina. This study characterized the potential source and the impacts of the outbreak in 2017–2019. Transect surveys indicated a sharp decrease in the density of live vicuña and guanaco by 68% and 77%, respectively, from May 2017 to June 2018. By April 2019 no vicuña or guanaco were recorded on transect surveys, suggesting a near-extinction at the local level. Clinical signs consistent with mange (e.g. intense pruritus, hyperkeratosis, alopecia) were observed in 24% of live vicuña (n = 478) and 33% of live guanaco (n = 12) during surveys, as well as in 94% of vicuña carcasses (n = 124) and 85% of guanaco carcasses (n = 20) opportunistically examined during the study period. Histological examination (n = 15) confirmed sarcoptic mange as the cause of the cutaneous lesions. Genetic characterization revealed that *Sarcoptes scabiei* recovered from seven vicuña (n = 13) and three guanaco (n = 11) shared the same genotype, which is consistent with a single source and recent origin of the epidemic. A governmental livestock incentive program introduced llama (*Lama glama*) in areas adjacent to SGNP in 2009, some of which reportedly had alopecic scaling consistent with sarcoptic mange. We hypothesize that the introduction of mange-infected llama may have triggered the outbreak in wild camelids which has now put them at a high risk of local extinction. This unprecedented event highlights that the accidental introduction of disease may be underestimated at the onset yet can have devastating effects on native ungulate populations with potentially profound effects at the community and ecosystem levels.

## Introduction

Emerging infectious diseases are caused by new pathogens or known pathogens that have recently increased their incidence or geographic distribution or have spread to new hosts [1]. In wild animals, such diseases have caused dramatic population declines, often leading to collapse and local extinction [1–5]. An emerging disease of increasing relevance for wildlife is sarcoptic mange, a highly contagious skin disease of mammals caused by the mite *Sarcoptes scabiei*.

Sarcoptic mange has been reported in at least 12 orders, 39 families and 148 species of domestic and wild mammals [6]. *S. scabiei* induces skin hypersensitivity, inflammation, intense itching, pain, and hair loss [7, 8]. As the disease progresses, the skin thickens and develops deep fissures, impairing thermoregulation. In seasonal climates, this may lead to a negative energy balance, progressive emaciation, and limited ability to forage and evade predators [9, 10].

The epidemiology of sarcoptic mange can vary considerably in wild animals depending on the geographic region and host species [7, 8]. While in some regions and species sarcoptic mange is endemic [11], in other regions it can cause devastating epidemics, leading to drastic population declines [12, 13]. Some outbreaks have been linked to transmission between livestock and wildlife [11]. Lavín et al (2000) [14] were able to reproduce the disease experimentally in the Cantabrian chamois (*Rupicapra pyrenaica parva*) from mites recovered from domestic goats (*Capra aegagrus hircus*), supporting the hypothesis that domestic animals may serve as a source of infection. There is also circumstantial evidence that the mange outbreaks that decimated Spanish ibex (*Capra pyrenaica hispanica*) likewise originated from domestic goats [12].

Sarcoptic mange is known to occur in wild South American camelids, vicuña (*Vicugna vicugna*) and guanaco (*Lama guanicoe*) [15–18]. However, information on the distribution and prevalence of this disease is extremely scarce, restricted to a low number of sites, and mostly reported in gray literature. In Argentina, both wild camelid species have suffered considerable retractions in their distribution and abundance due to hunting, competition with livestock, and habitat loss, resulting in reductions of up to 40% in the original distribution of guanaco and the near-extinction of vicuña in the mid-twentieth century [19]. Currently, guanaco and vicuña are globally categorized as “Least Concern”, though some sub-populations are small, fragmented, and isolated, rendering them locally susceptible to stochastic events [20, 21].

San Guillermo National Park (SGNP) protects the largest sympatric vicuña and guanaco populations in Argentina and represents the southern limit of the distribution of vicuña [22]. In 2014, sarcoptic mange was diagnosed for the first time in both camelid species at SGNP. Soon thereafter, significant declines in the camelid populations were noticed as the number of affected animals increased [23]. The abrupt nature and rapid progression of the outbreak suggests a recent introduction of the causative pathogen [24, 25].

A governmental livestock incentive program introduced llama (*Lama glama*) in areas adjacent to SGNP in 2009 [26], some of which were diagnosed with mange, suggesting a potential link between the two events. Here, we explore the hypothesis of llama-sourced infection by describing epidemiological aspects of the outbreak during 2017-19, genetically characterizing the mites, and analyzing the spatial and temporal overlap of the events.

## Materials and Methods

### Study area

San Guillermo National Park (SGNP) is in San Juan province, Argentina (29°4’12”S, 69°21’0”W) (Fig 1) and covers 166,000 ha between 2,000 and 5,600 meters above sea level. San Guillermo Provincial Reserve (SGPR) (about 815,460 ha) surrounds SGNP to the northwest. Together, SGNP and SGPR make up San Guillermo Biosphere Reserve (SGBR) (Fig 1), which protects 981,460 ha of the Puna and High Andes eco-regions [22].

**Figure 1:**
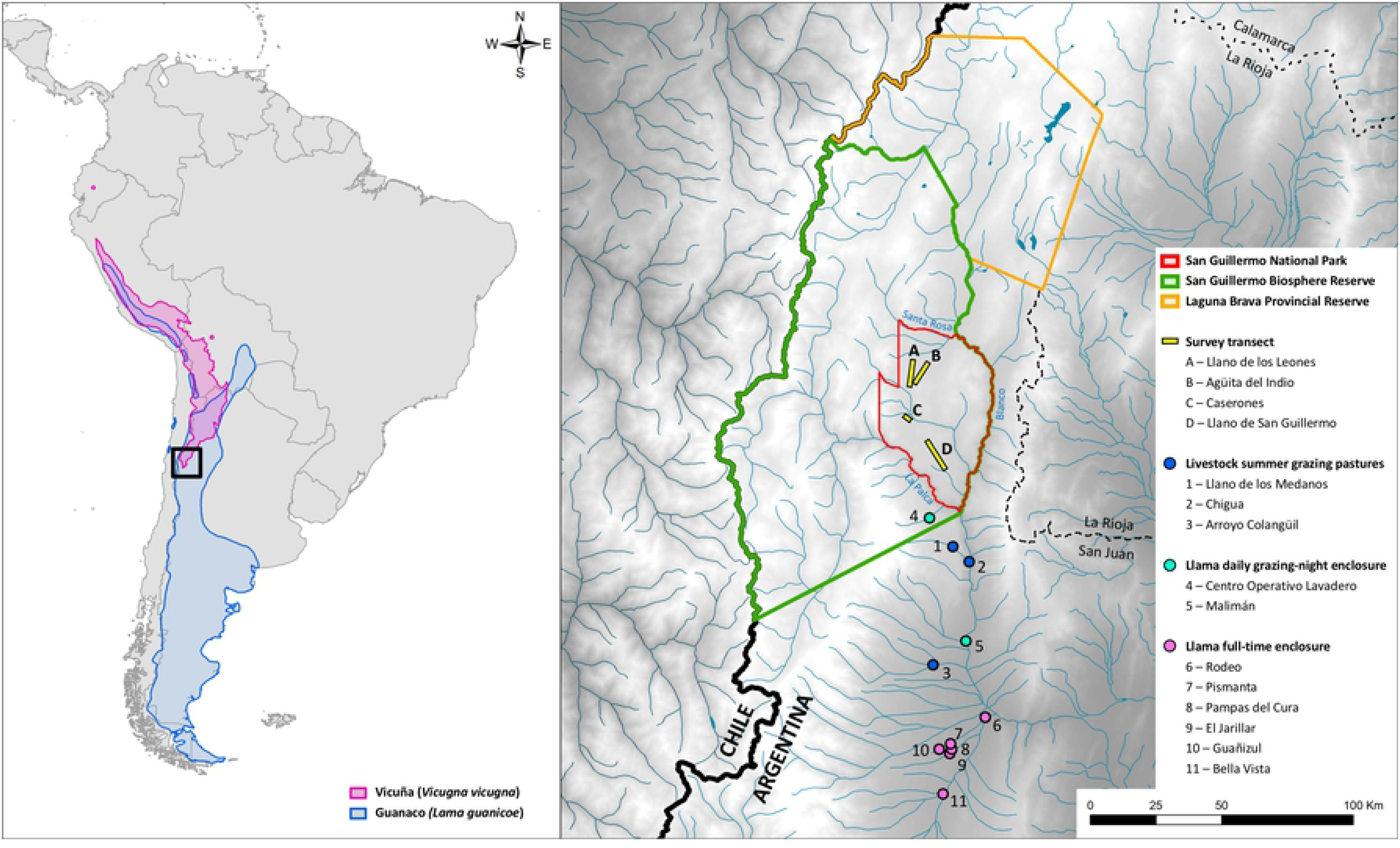
Location of San Guillermo National Park and study transects in relation to the natural distribution of vicuña (*Vicugna vicugna*) and guanaco (*Lama guanicoe*) and nearby farms and grazing areas of livestock and recently introduced llama (*Lama glama*).

Rainfall is scarce with an average 240 mm per year, concentrated in January-March. The temperature range is −23 °C to 27 °C; January is the warmest (average 14 °C) and July the coldest month (average −1 °C) [27]. SGNP is characterized by extensive open plains (81% of the study area) located at 3,400 meters above sea level, surrounded by hills and mountain peaks. These plains are traversed by narrow canyons (10-300 m wide) with steep rocky walls representing 15% of the study area, and a few isolated flooded meadows within the plains or on the riverbanks account for the remaining 4% of the study area [28]. Vicuña are ten times more abundant than guanaco in the park [29].

Vicuña and guanaco concentrate in high altitude meadows and plains [30], but guanaco migrate during the winter to low altitude areas, where they are exposed to goats and sheep and to a lesser extent, cattle [30]. In 2009, some farmers nearby SGNP received llamas (*Lama glama*) through a San Juan province governmental livestock incentive program (‘Programa Camélidos de los Andes’) [26]. A group of these animals were temporarily housed within SGPR, at Centro Operativo Lavadero (Fig 1, site 4).

### Population density and proportion of live individuals infected with mange

#### Density

After the onset of the sarcoptic mange outbreak in 2014, increasing prevalence of mange lesions and death of wild camelids were recorded in SGNP [23]. To estimate the density of remaining live vicuña and guanaco, in our study we conducted five field surveys in May, September and December 2017, and April and June 2018. In each survey we implemented four transects following park dirt roads: Llano de los Leones, Llano San Guillermo, Caserones and Agüita del Indio (Fig 1; further details provided in Table S1). We used binoculars (Tasco® 7 x 35 mm) and a telescope (Bushnell® 15-45 x 60 mm) to search for and observe animals and laser rangefinders (Bushnell® Elite 1500 and Bushnell® DX 1000) to measure distances between the animals and the observer. We traveled transects at a speed of 20 km/h. When we spotted animals, we stopped and observed them for 5-10 min to count them and confirm infection status. We travelled the transects twice at one day intervals and animals observed within 1,000 m on each side of the transect were counted. Along each transect, we recorded the distance from and the angle of the animal or group of animals to the transect. Densities were estimated using Distance 7.1 [31]. Due to small sample sizes, data were grouped by species in three periods covered by our study (May 2017, September-December 2017 and April-June 2018) and no estimates could be obtained for September 2018 and April 2019. The probability of detection was calculated for each period and species, provided it reached a minimum of 60-80 observations [31]. Thus, for vicuña, detection probabilities were obtained for each of the three periods, while for guanaco they were obtained by grouping the data with those of vicuña due to low counts.

### Mange infection

To quantify the proportion of live camelids with mange we conducted eight field surveys in February, May, September, and December 2017; April, June, and September 2018; and April 2019 (Table S1). We used the same transects but only evaluated individuals within 200 m from the transect because of the unfeasibility of accurately identifying infected animals at distances ≥ 200 m. Because incipient infections are undetectable without handling the animals, our estimates represent the minimum infected proportion. We identified cases of sarcoptic mange when one or more of the following signs were observed: intense scratching, difficulty walking, thickening, crusty or cracked skin, and alopecia or ruffled or detached fiber. According to the clinical stage of disease, we used three categories: (A) early stage, only pruritus evident; (B) advanced stage, difficulty walking and/or visible injuries to the limbs; and (C) severe stage (Fig 2 A, B and C), alopecia extending to several parts of the body. Because the categories represent increasing severity, each level includes the signs of the previous one. To avoid inter-observer bias, all observations were made by the same person (H. Ferreyra).

**Figure 2:**
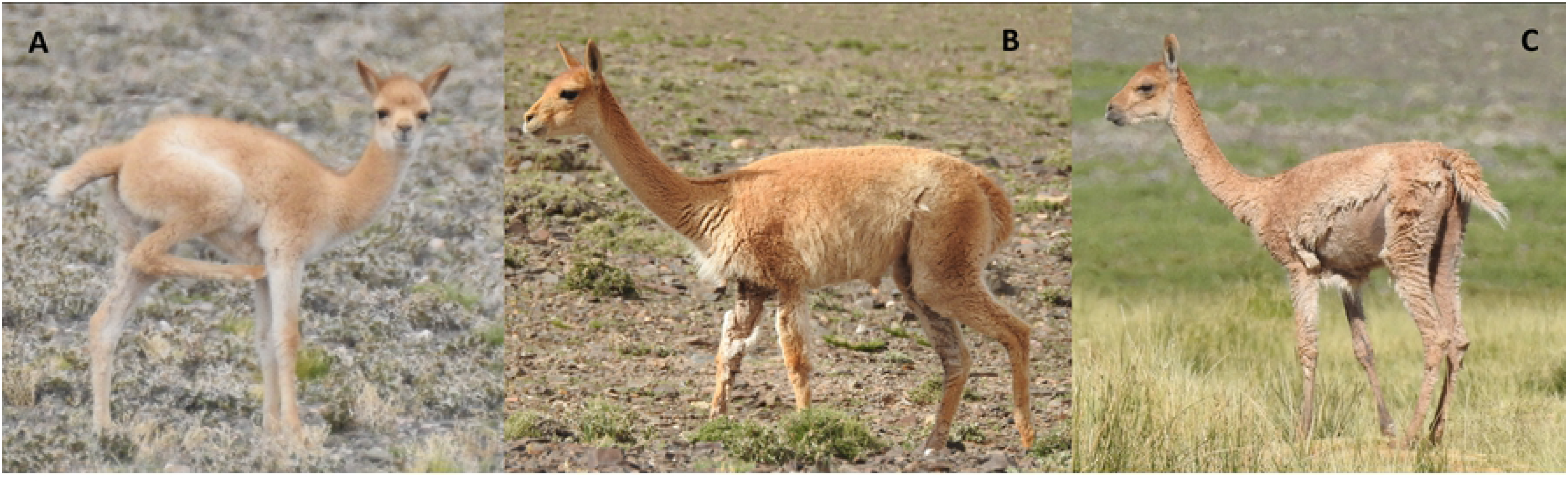
Clinical stage of mange in live vicuña: (A) early stage, (B) advanced stage and (C) severe stage.

### Mange infection in carcasses and mite collection

Camelid carcasses found during opportunistic searches throughout SGNP from May 2017 to June 2018 were evaluated for the presence of skin lesions consistent with mange. Carcasses were recorded only if the remains included limbs with skin and skull. Examined carcasses were identified with a cattle marker pen, and the lower jaw was disarticulated to avoid double counts between surveys. Carcasses were classified as either mange positive or negative (severity was not determined), and when possible, lesions were scraped with a sterile surgical blade. Scrapings were preserved in two vials, one with mineral oil for microscopy and the other with 70% ethanol for genotyping [32]. *S. scabiei* was confirmed microscopically following standard guidelines [33]. Using a magnifying glass and watchmaker’s tweezers, individual mites were recovered and placed in 70% ethanol.

Affected skin samples from fresh, whole carcasses, were also preserved in 10% formalin for histopathology. In these cases, body condition was estimated as good, poor and emaciated according to [34] (2009). Tooth wear and replacement from the lower jaw teeth were used to determine age class (cria, juvenile and adult) [35].

### Tracing the outbreak source

Based on the hypothesis that introduced llama were linked to the outbreak in wild camelids, we interviewed the veterinarians who worked in the governmental livestock incentive program that introduced llama to the study area. Interviewees were asked about the number, geographic origin, dates of arrival and health care and husbandry of the introduced llama over the duration of the program. Additional information included where llamas were initially housed, and whether mange was noted at any time. Likewise, llama breeders in proximity to the park were visited (Fig 1) and information was compiled on the current number of animals in their herds, whether they were corralled or allowed to graze freely during the day and confined at night, whether they had noticed signs of mange, and if they received clinical exams and preventive treatment for mange.

To assess whether mange had been historically detected in wild camelids in the area we posed the question “do you remember seeing or have you heard about mange in wildlife in this area in the last fifty years” to provincial and national park rangers, as well as to technicians from the Secretary of the Environment of San Juan province. The latter then extended the question to elderly farmers (˃60 years) in the region when they were visited for other purposes.

### Statistical analysis

The proportion of mange-infected individuals and their respective 95% confidence intervals (CI) were determined for both live and dead camelids. Because the abundance of guanaco is low at SGNP, further statistical analysis were conducted only for vicuña. Likelihood ratio chi-square tests were used to evaluate whether the proportion of vicuña with mange or the proportion of disease clinical stage categories were unevenly distributed relative to age classes (excluding “not determined”), transects and survey months. Binary logistic regression was used to evaluate whether age classes, transects and months were significant predictors of the proportion of live vicuña with mange. Multinominal logistic regression was used to evaluate whether age classes (reference category = adult; excluding “not determined”), transects (ref = Agüita del Indio) and survey months (ref = April 2018) were significant predictors of the different stages of mange (early, advanced, severe; reference category = without mange) of live vicuña. Odds ratios (OR) and their 95% confidence intervals were calculated for pairwise comparisons among categories of variables identified as significant (i.e. those where the OR CI interval did not include 1).

### Genetic characterization of mites

The Micro DNA Extraction Kit (Qiagen, Valencia, CA) procedure was used for the preparation of mite DNA from a single *Sarcoptes* mite sample according to the manufacturer’s recommendations. Prior to individual DNA extraction, dead mites were pierced with an 18-gauge needle under a dissecting microscope and digested overnight in lysis buffer and proteinase K at 56 °C (Qiagen, Valencia, CA). Final DNA product from each mite was eluted in 60 µL of AE buffer. We selected 10 published microsatellite markers (SARMS 33-38, 40, 41, 44, and 45) [36] to genotype individual mites. Forward primers were labeled with HEX or 6-FAM dye (Integrated DNA Technologies, Coralville, IA) and reconstituted into 100 µM working dilutions. Primer pairs were combined into paired multiplex with 1.5 – 2.5 µM of each primer. We performed PCR using the Qiagen 2X Type-it Multiplex PCR Master Mix, 10X multiplex primer mix (2.5 µL), DNA-free water (7 µL), and 2-3 µL DNA for a total reaction of 25 µL. Thermocycling conditions were as published [36]. PCR products were transferred to 96-well plates (Biotix Inc, San Diego, CA) for electrophoresis and digital measurement of length polymorphisms on an ABI 3730 analyzer (Perkin-Elmer Davis, CA) using the program STRand (Veterinary Genetics Laboratory, University of California, Davis, CA;[37]). Microsatellite scoring and allele binning were performed with the R-package MsatAllele [38].

Data was reformatted using CREATE v1.37 [39], and descriptive statistics and diversity analyses were carried out with GenAlEx v. 6.2 [40], ML-Relate [41], and R package [42] (R Core Development Team 2018) PopGenReport [43] and poppr [44] to determine the number of private alleles, allele frequencies and expected (He) and observed (Ho) heterozygosity, and also to test for Hardy–Weinberg equilibrium (HWE), and partitioned components of variance using analysis of molecular variance (AMOVA). To evaluate differentiation among the *S. scabiei* mite populations, we calculated the pairwise F-statistic. Possible errors in genotyping due to stuttering of large allele dropout were evaluated using MicroChecker v.2.2.0.3 [45]. Null alleles were estimated using ML-Relate. P-values ≤ 0.05 were considered statistically significant.

## Results

### Field data

From May 2017 to June 2018, the population density declined from 8.89 to 2.87 individuals/km^2^ for vicuña (68% decrease) and from 0.26 to 0.06 individuals/km^2^ for guanaco (77% decrease) at SGNP (Table 1). Figure 3 summarizes the temporal distribution of the number of camelids with and without mange recorded during transect surveys (live individuals) or opportunistically recorded (dead individuals). No guanaco were seen during mange-detection transect surveys (≥ 200 m on both sides of transect) in June 2018, September 2018 and April 2019, and no vicuña were seen during these surveys in April 2019.

**Table 1:**
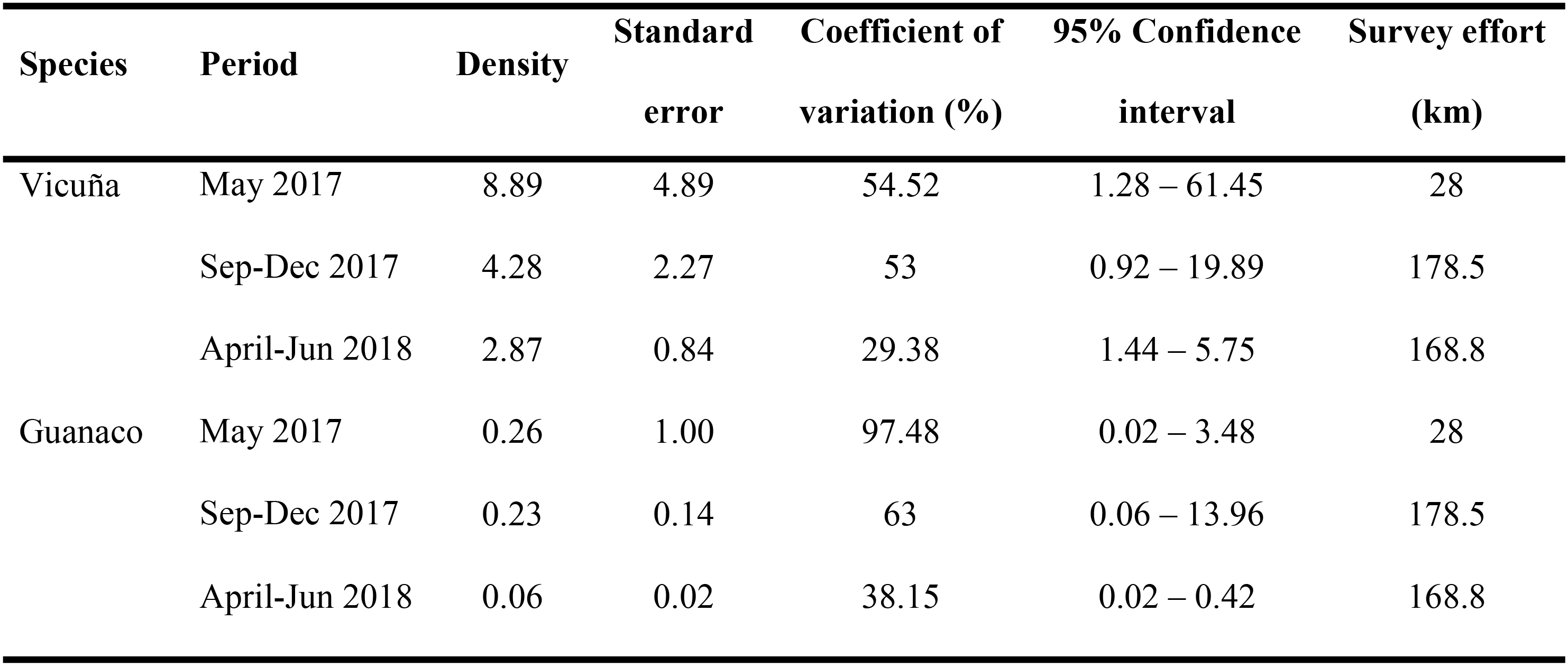
Density (individuals/km^2^) of vicuña and guanaco at San Guillermo National Park, May 2017 – June 2018.

**Figure 3:**
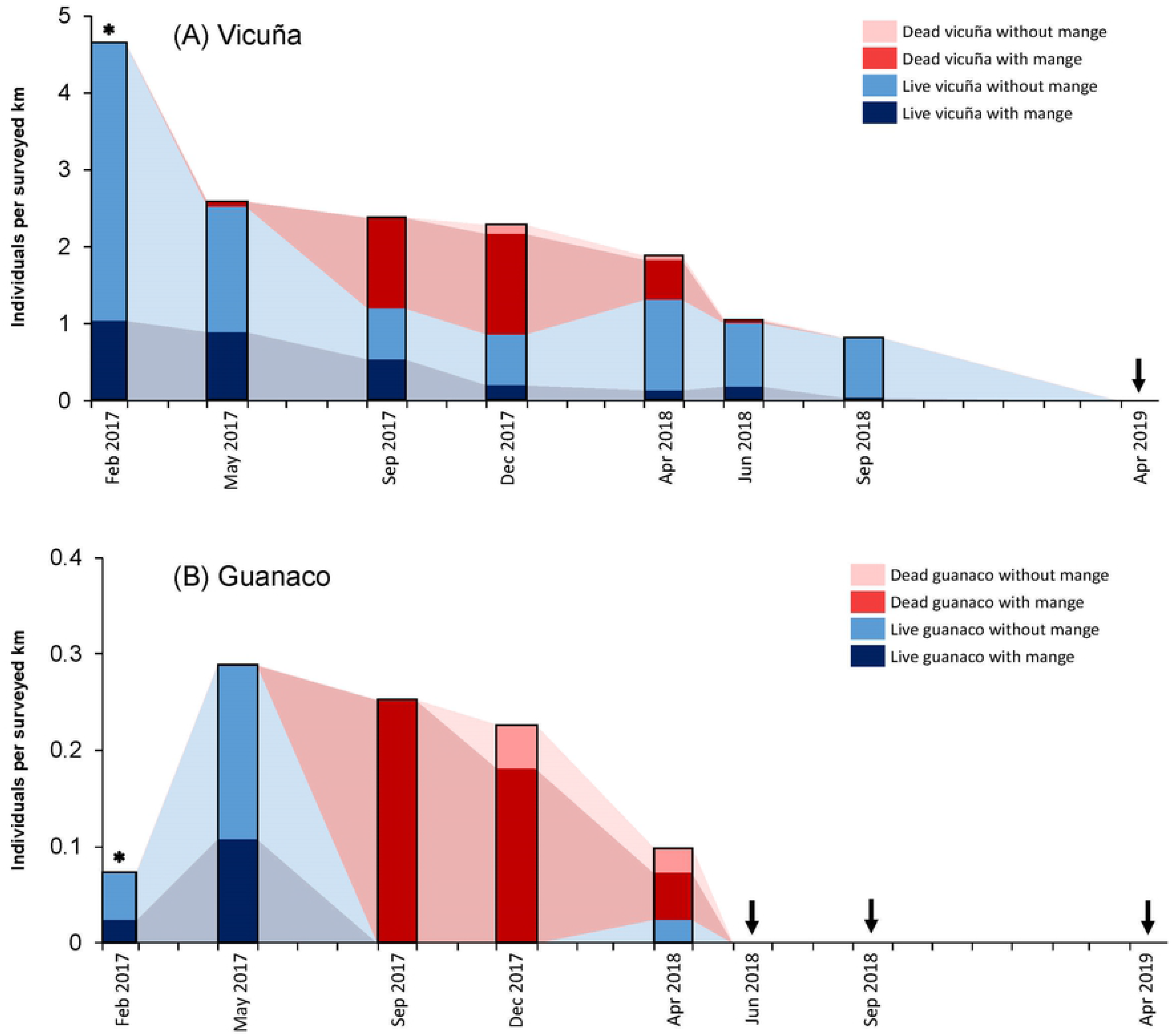
Time series of the number of live camelids observed in transect surveys and opportunistically collected carcasses with and without mange at San Guillermo National Park, February 2017 – April 2019. Asterisks indicate months when carcasses were not evaluated. Arrows represent field surveys where no individuals were recorded. Light shaded areas between bars are used to highlight the relative changes between field surveys (no data was collected in these intervals).

During the study period, 24.1% (CI = 20.3 – 28.2; n = 478) of live vicuña met our case definition for mange (Table 2). Only twelve live guanaco were seen during transect surveys: (a) three individuals in February 2017 (two adults without mange at Llano San Guillermo, one adult with mange at Caserones), (b) eight individuals at Llano San Guillermo in May 2017 (one cria without mange, three adults with mange, four individuals of unknown age group without mange), and (c) one individual at Caserones in April 2018 (without mange). The prevalence of mange in live guanaco was therefore 33.3% (CI = 9.9 – 65.1; n = 12), and all cases were in the advanced-stage category.

**Table 2:** Number and proportion of live vicuña with mange recorded during transect surveys at San Guillermo National Park, February 2017 – April 2019. The stages of clinical disease were categorized as: (A) early stage, only pruritus evident, (B) advanced stage, difficulty walking and/or visible injuries to the limbs, and (C) severe stage, alopecia extending to several parts of the body. Because the categories represent increasing severity, each level includes the signs of the previous one.

|  | Individuals<br>examined | Individuals<br>with mange | Proportion<br>(95% CI) | Early<br>stage | Advanced<br>stage | Severe<br>stage |
| --- | --- | --- | --- | --- | --- | --- |
| *Age class* |  |  |  |  |  |  |
| Cria | 86 | 13 | 15.1% (8.3 – 24.5) | 10 (83%) | 2 (17%) | 0 |
| Juvenile | 137 | 37 | 27% (19.8 – 35.3) | 2 (5%) | 22 (59%) | 13 (35%) |
| Adult | 205 | 59 | 28.8% (22.7 – 35.5) | 6 (11%) | 35 (65%) | 13 (24%) |
| Not determined | 50 | 6 | 12% (4.5 – 24.3) | 1 (8%) | 9 (75%) | 2 (17%) |
| *Month* |  |  |  |  |  |  |
| February 2017 | 189 | 42 | 22.2% (16.5 – 28.8) | 7 (17%) | 19 (45%) | 16 (38%) |
| May 2017 | 70 | 25 | 35.7% (24.6 – 48.1) | 7 (28%) | 16 (64%) | 2 (8%) |
| September 2017 | 53 | 24 | 45.3% (31.6 – 59.6) | 1 (4%) | 18 (75%) | 5 (21%) |
| December 2017 | 35 | 8 | 22.9% (10.4 – 40.1) | 2 (25%) | 4 (50%) | 2 (25%) |
| April 2018 | 54 | 6 | 11.1% (4.2 – 22.6) | 0 | 4 (67%) | 2 (33%) |
| June 2018 | 43 | 8 | 18.6% (8.4 – 33.4) | 2 (25%) | 5 (63%) | 1 (13%) |
| September 2018 | 34 | 2 | 5.9% (0.7 – 19.7) | 0 | 2 (100%) | 0 |
| April 2019 | 0 |  |  |  |  |  |
| *Transect* |  |  |  |  |  |  |
| Llano de los Leones | 159 | 54 | 34% (26.7 – 41.9) | 10 (19%) | 28 (53%) | 15 (28%) |
| Agüita del Indio | 184 | 28 | 15.2% (10.4 – 21.2) | 1 (3%) | 21 (72%) | 7 (24%) |
| Caserones | 15 | 4 | 26.7% (7.8 – 55.1) | 0 | 4 (100%) | 0 |
| Llano San Guillermo | 120 | 29 | 24.2% (16.8 – 32.8) | 8 (28%) | 15 (52%) | 6 (21%) |
| **Total** | **478** | **115** | **24.1% (20.3 – 28.2)** | **19 (17%)** | **68 (59%)** | **28 (24%)** |

The overall proportion of live individuals with mange was similar in vicuña and guanaco (LRT = 0.511, df = 1, P = 0.475). The proportion of live vicuña with mange was significantly different between survey months (LRT = 31.72, df = 6, P < 0.001), transects (LRT = 16.65, df = 3, P = 0.001) and age classes (LRT = 6.747, df = 2, P = 0.034). Binary logistic regression indicated that only the survey month (P = 0.001) and transect (P < 0.001) were significant predictors of the proportion of live vicuña with mange, whereas age class was not (P = 0.163). Specifically, the pairwise comparisons revealed that live vicuña recorded in September 2017 were more likely to have mange than those recorded in February 2017 (OR = 4.02), April 2018 (OR = 11.26) and September 2018 (OR = 9.09) (95% CIs provided in Table S2). Additionally, live vicuña recorded at Caserones (OR = 5.77), Llano de los Leones (OR = 4.59), and Llano San Guillermo (OR = 3.05) were more likely to have mange than those recorded at Agüita del Indio (95% CIs provided in Table S2).

Among live vicuña with mange, the proportion of individuals in each disease stage category varied significantly relative to the survey months (LRT = 50.77, df = 18, P < 0.001), transects (LRT = 22.81, df = 6, P < 0.001; “Caserones” omitted) and age classes (LRT = 40.45, df = 6, P < 0.001) (Fig 2). Multinominal logistic regression for live vicuña revealed that: (a) cria were more likely to present early stage disease (OR = 4.86) and less likely to present advanced stage disease (OR=0.14) relative to adults; (b) individuals recorded in February 2017 were more likely to present advanced stage disease than those recorded in May 2017 (OR = 3.16), September 2017 (OR = 11.02) and June 2018 (OR = 7.85); and (c) individuals recorded at Agüita del Indio were less likely to present early stage disease than those recorded at Llano de los Leones (OR = 0.08) and Llano San Guillermo (OR = 0.06), less likely to present advanced stage disease than those recorded at Caserones (OR = 0.08) and Llano de los Leones (OR = 0.22) and less likely to present severe stage disease than those recorded at Llano de los Leones (OR = 0.18) (95% CIs provided in Table S3).

Among opportunistically-collected carcasses, 93.5% of vicuña (CI = 87.7 – 97.2; n = 124) and 85.0% of guanaco (CI = 62.1 – 96.8, n = 20) met our case definition (Table 2). The overall proportion of dead individuals with mange was similar in vicuña and guanaco (LRT = 1.485, df = 1, P = 0.223). The proportion of dead vicuña with mange was similar among survey months (LRT = 4.725, df = 2, P = 0.094; “May 2017” and “June 2018” omitted) and age classes (LRT = 5.682, df = 2, P = 0.058) (Table 3). During carcass searches, only five animals (four vicuña and one guanaco), were found whole and fresh (recently predated by cougar *Puma concolor*). These carcasses presented with advanced clinical stage of mange, and all were in good body condition with good musculature.

**Table 3:** Number and proportion of mange in examined vicuña and guanaco carcasses at San Guillermo National Park, May 2017 – June 2018.

| Category | Vicuña |  |  | Guanaco |  |  |
| --- | --- | --- | --- | --- | --- | --- |
|  | Individuals<br>examined | Individuals<br>with mange | Proportion<br>(95% CI) | Individuals<br>examined | Individuals<br>with mange | Proportion<br>(95% CI) |
| *Age class* |  |  |  |  |  |  |
| Cria | 23 | 19 | 82.6% (61.2 – 95.1) | 1 | 1 | 100% (2.5 – 100) |
| Juvenile | 22 | 20 | 90.9% (70.8 – 98.9) | 2 | 2 | 100% (15.8 – 100) |
| Adult | 75 | 73 | 97.3% (90.7 – 99.7) | 15 | 13 | 86.7% (59.5 – 98.3) |
| Not determined | 4 | 4 | 100% (39.8 – 100) | 2 | 1 | 50% (1.3 – 98.7) |
| *Month* |  |  |  |  |  |  |
| May 2017 | 3 | 3 | 100% (29.2 – 100) | 0 |  |  |
| September 2017 | 33 | 33 | 100% (89.4 – 100) | 7 | 7 | 100% (59 – 100) |
| December 2017 | 63 | 58 | 92.1% (82.4 – 97.4) | 10 | 8 | 80% (44.4 – 97.5) |
| April 2018 | 23 | 21 | 91.3% (72 – 98.9) | 3 | 2 | 66.7% (9.4 – 99.2) |
| June 2018 | 2 | 1 | 50% (1.3 – 98.7) | 0 |  |  |
| **Total** | 124 | 116 | 93.5% (87.7 – 97.2) | 20 | 17 | 85% (62.1 – 96.8) |

### Histological findings

Histology from 14 vicuña and one guanaco carcasses revealed typical sarcoptic mange lesions with abundant mites in all specimens. Histological findings were consistent with chronicity such as hyperplasia of the epidermis and of sebaceous glands (15/15), collagen sclerosis (12/15), as well as acute changes like presence of inflammatory cells (neutrophils, eosinophils) and congested blood vessels in all cases (15/15, 100%) (Fig S1 A, B, C and D). Lesions identified as chronic histologically were more common in the axillary and inguinal regions of the body and coincided with areas where the greatest thickening of skin with crusts were observed macroscopically (Fig 4 A, B and C).

**Figure 4:**
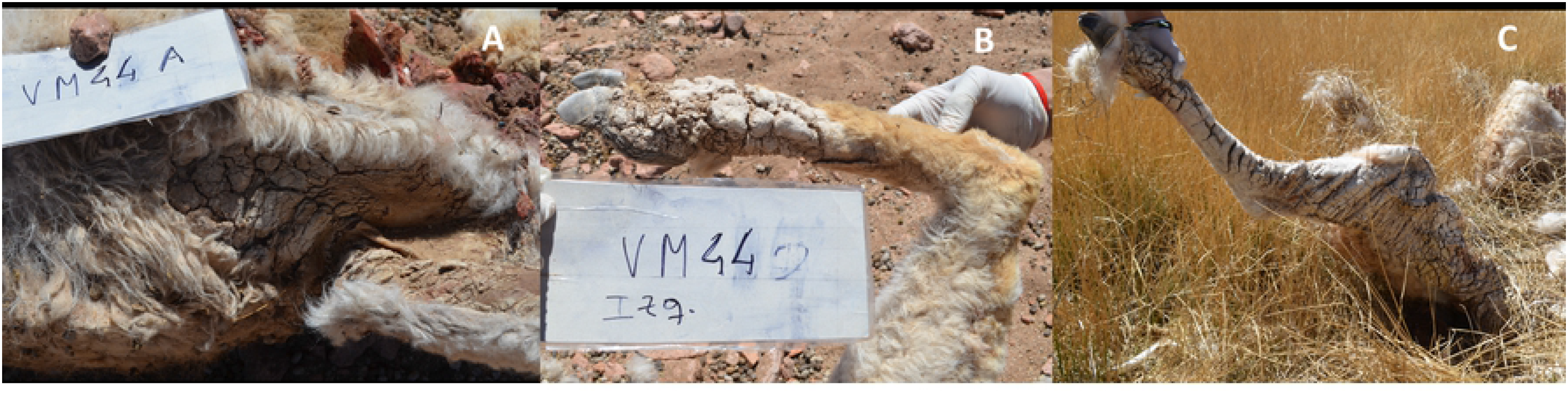
(A) vicuña carcass: scabs and deep cracks in the axillary area; (B) vicuña carcass: scabs and deep cracks on hind limb; (C) guanaco carcass: scabs and deep cracks along the hind limb and groin.

### Tracing the outbreak source

Five veterinarians were interviewed. Four participated in the livestock incentive program (‘Camélidos de los Andes’) between 2009 and 2011. A fifth veterinarian runs a large-animal practice in the town of Rodeo, near SGNP (see Fig. 1). According to their records, 156 llama entered San Juan province between 2009-2011 from Jujuy and Catamarca provinces (900 and 300 km north of San Juan, respectively). Veterinarians reported that llama were initially confined to a community pen in Rodeo where mange was detected in at least two animals upon arrival from Jujuy province in 2009 (Fig S2 A and B). Infected llama were treated with ivermectin. Llama were given to local farmers between 2009-2011. About 15 llama that were not claimed by farmers were placed under temporary care of provincial park rangers at a ranger post, Centro Operativo Lavadero (Lavadero), within SGPR.

During our study period, seven facilities that housed llama were identified. All were private farms in Iglesias department (Fig 1). The farmer in Malimán (since 2009) and the rangers in Lavadero (mentioned above) allowed llama to graze freely. Llama in Lavadero were reportedly moving about 8 km to the northwest into SGNP on a daily basis, and in Malimán, their space use overlapped with that of guanaco. According to two interviewees, no mange was observed at these two sites, although there was no sustained veterinary care due to the expiration of the government program in 2013-2014.

Finally, the extended interviews revealed that in the last 20 years, there were no outbreaks of mange in non-camelid livestock in proximity of SGNP or SGBR. Likewise, there were no previous reports of sarcoptic mange outbreaks in wild camelids in the area at least in the last 50 years since SGPR was established.

### Mite characterization

A total of 24 mites were selected for molecular identification from the skin scrapings of vicuña and guanaco; 13 mites collected from seven vicuña and 11 mites collected from three guanaco. Sixteen alleles were detected across 10 microsatellite loci. Depending on the loci, allele count ranged from one (SARM-36 and 38) to three (SARM-33 and 40). A total of 6 private alleles (i.e. alleles found only in one population and among the broader collective populations of study) were detected and distributed among eight loci (SARM-33, 35, 37, and 40). The distribution and allele frequencies among populations of *S. scabiei* mites according to the host is presented in Table 4.

**Table 4:** Frequency of alleles by population. Distributions of allele frequencies in 10 microsatellite loci among *Sarcoptes scabiei* mite populations by host, vicuña and guanaco (allele sizes are in base pairs). N is the number of mites collected and genotyped from seven vicuña and three guanaco at each allele. Private alleles are denoted with “^†^”.

| Locus | Allele | Vicuña Mites | Guanaco Mites |
| --- | --- | --- | --- |
| SARM-33 | N | 12 | 11 |
|  | 245 | 0.083<sup>†</sup> | 0.000 |
|  | 247 | 0.875 | 1.000 |
|  | 274 | 0.042<sup>†</sup> | 0.000 |
| SARM-45 | N | 13 | 11 |
|  | 194 | 1.000 | 1.000 |
| SARM-35 | N | 12 | 11 |
|  | 136 | 1.000 | 0.818 |
|  | 138 | 0.000 | 0.182<sup>†</sup> |
| SARM-38 | N | 13 | 11 |
|  | 211 | 1.000 | 1.000 |
| SARM-34 | N | 13 | 11 |
|  | 209 | 1.000 | 1.000 |
| SARM-44 | N | 13 | 11 |
|  | 270 | 1.000 | 0.909 |
|  | 272 | 0.000 | 0.091<sup>†</sup> |
| SARM-40 | N | 13 | 11 |
|  | 248 | 0.154<sup>†</sup> | 0.000 |
|  | 250 | 0.846 | 1.000 |
| SARM-41 | N | 13 | 11 |
|  | 236 | 1.000 | 1.000 |
| SARM-36 | N | 13 | 11 |
|  | 272 | 1.000 | 1.000 |

| **SARM-37** | **N** | **13** | **11** |
| --- | --- | --- | --- |
|  | **180** | 0.923 | 1.000 |
|  | **274** | 0.077<sup>†</sup> | 0.000 |

Vicuña-derived mites had more total alleles detected overall (n = 14) compared to mites collected from guanaco (n = 12); however, both mite-derived populations displayed relatively low allelic richness (R_vicuña_ = 1.35, R_guanaco_ = 1.19, Table 5). Further, mites from vicuña and guanaco presented relatively few alleles with a low occurrence of polymorphisms, 30% polymorphic loci in vicuña-derived mites and 20% in guanaco-derived mites. Fixed alleles were detected for both vicuña and guanaco-derived mites at SARM-34, 36, 38, 41, and 45 (Table S4). Fixed alleles were also observed for vicuña- derived mites at SARM-35, 37, and 44, while additional fixed alleles for guanaco-derived mites were detected at SARM-33 and 40. Values of expected (He) and observed (H_o_) heterozygosity were also low for mites collected from vicuña (H_e_ = 0.063, H_o_ = 0.024) and guanaco (H_e_ = 0.046, H_o_ = 0.055).

**Table 5:**
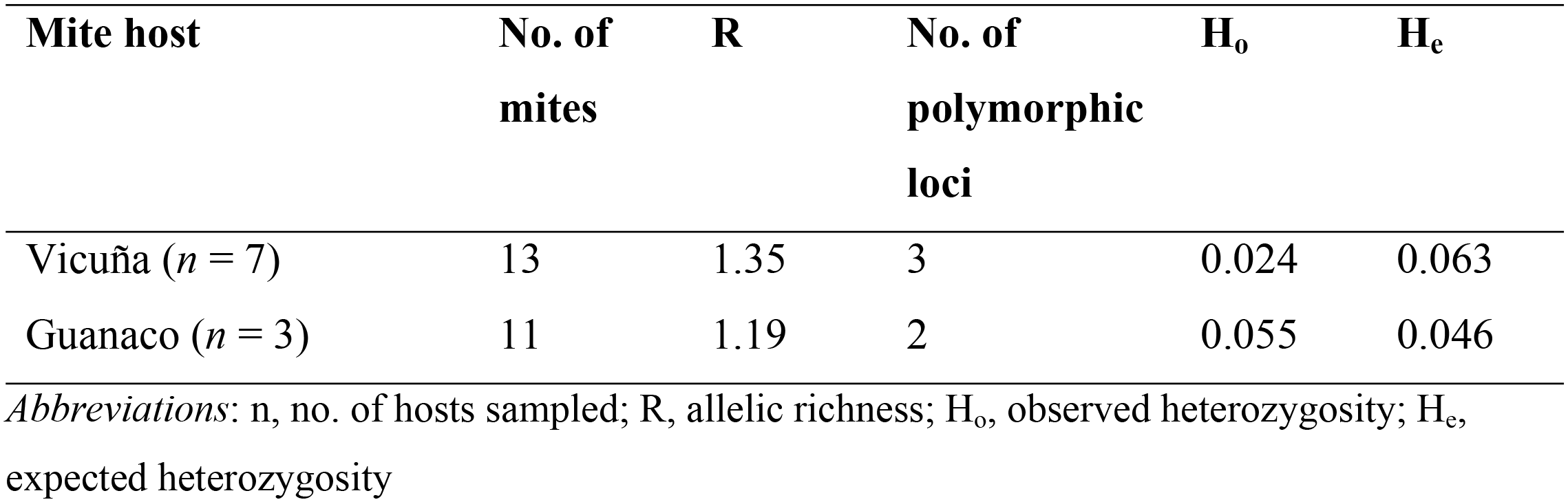
Characteristics of genetic variability of *Sarcoptes scabiei* obtained from vicuña and guanaco carcasses in San Guillermo National Park.

| **Mite host** | **No. of mites** | **R** | **No. of polymorphic loci** | **$H_o$** | **$H_e$** |
| --- | --- | --- | --- | --- | --- |
| Vicuña ($n = 7$) | 13 | 1.35 | 3 | 0.024 | 0.063 |
| Guanaco ($n = 3$) | 11 | 1.19 | 2 | 0.055 | 0.046 |

*Abbreviations:* n, no. of hosts sampled; R, allelic richness; $H_o$ , observed heterozygosity; $H_e$ , expected heterozygosity

Mites from guanaco showed no significant deviations from the Hardy-Weinberg equilibrium (HWE), while mites from vicuña had significant HWE departures at SARM-33 (P = 0.032) and SARM-40 (P = 0.004). AMOVA analysis showed the highest percentage of variance to occur within samples (57.7%, P = 0.04) rather than between populations (6.35%, P = 0.01). Pairwise F_ST_ values demonstrated both populations were closely related (F_ST_ = 0.054, P = 0.025).

## Discussion

Sarcoptic mange is an emerging global wildlife disease. Recent reported cases worldwide reflect broad geographic spread, an increase in host species and greater virulence, and have been associated with population declines [6]. Here we report an outbreak of mange with a devastating effect on wild camelid populations within a protected area and its potential link with introduction of llama in the vicinity.

The impact of mange on the abundance of wild camelids in SGNP was severe. This study, which spanned a period of 26 months (February 2017 – April 2019), took place at an advanced stage of the epidemic, when the population reduction was most drastic. A decline in population densities of 67 and 77% for vicuña and guanaco, respectively, between May 2017 and June 2018, coupled with the continuous decrease in individual counts through April 2019, reflect the near disappearance of these species from the park by the end of this study (Tables 1 and 2, Fig 3).

Despite the continuous numerical decrease of camelids in the park, mange persisted at the end of this study, albeit at low rates. By April 2019, only one mange-infected vicuña in advanced clinical stage was found by doubling the length (24 km) of the Agüita del Indio transect (data not shown). This suggests that mechanisms independent of density were involved in transmission, such as frequency-dependent mechanisms (e.g. mating behavior), that allow a pathogen to continue to spread even when population size declines to the point of near local extinction [46, 47]. Indirect transmission through contact with contaminated objects [46, 48, 49] is also possible. In particular, the role of communal sites such as dust baths or other elements of the environment like shrubs (in this study it was observed that animals used hard vegetation to scratch) in the transmission of mange remain unknown. The severe hyperkeratotic or crusted clinical form of mange observed in this study is characterized by high load of mites and is thus highly contagious [50].

Mange-infected camelids were seen throughout the study period. However, the proportion of live affected animals may have been underestimated due to limitations in detection of early stages from distant observation. Conversely, the proportion of mange-infected carcasses was high despite examination of mostly limbs with skin remains, which may have missed infection in other parts of the body. The occurrence of mange in live vicuña was similar across age classes, but severity varied, and severe stages were not observed in crias. Because crias were seen nursing from severely ill mothers, it is possible that lack of maternal care led to mortality of this age class before mange progressed. A higher proportion of vicuña in advanced stage of the disease, at which there is visible difficulty in their movements, was seen in Llano de los Leones. It is possible that mange influenced the distribution pattern of sick animals which congregated in a few flooded meadows, where food and water were easily accessible. These sites are also the preferred hunting sites for puma [51, 52], which may explain the steady numerical declines and the removal of animals before they reached severe stages of disease. Preliminary data show that the percentage of puma-killed camelids (n = 392) with mange lesions increased from 5 to 90% in 24 months at the outbreak onset (E. Donadio, unpublished data).

Spatially, the outbreak was initially detected in both SGNP and the larger SGPR. However, over time, infected camelids were observed in adjacent, outside park boundary areas. For example, mange-infected vicuña and guanaco were reported to the north of the park in 2016 (La Brava Reserve in La Rioja province, Fig 1) and infected guanaco were seen to the northeast, in San Juan province in 2018. While vicuñas are naturally restricted to high altitude locations, guanacos in this region are migratory and more prone to overlap with livestock, and recently, with introduced llama. The altitudinal migration of guanaco in the Andean Mountain range is a reported phenomenon [53, 54]. Moreover, home ranges of 1853 km^2^ (185,000 ha) have been described for guanaco in Argentina [55], showing the biological capacity of this species to move over large areas and their potential to disperse mange if infected. Guanaco could have thus acted as a bridge species for the transmission of the mite during its migration towards the high Andes inhabited by vicuña.

Interviewed veterinarians reported that the only cases of mange near SGNP in the last 20 years occurred in llama in 2009, when they were first introduced to San Juan province. Mange was specifically reported in a llama herd from Cieneguillas, Jujuy province, a site where mange is common in domestic and wild camelids [17]. The veterinarians reported that one of the infected llamas was treated for mange and then handed to a farmer in Malimán, who allowed his herd to graze freely and comingle with free-ranging guanaco.

This situation may have also occurred in the higher-altitude Lavadero, the ranger post adjacent to SGNP, where both vicuña and guanaco are present. The existence of more such sites of spatial overlap cannot be ruled out. From the extended interviews in San Juan, it is evident that at least in the last 50 years, mange had never been reported in wild camelids in the SGBR or its area of influence.

The guanaco and vicuña mites evaluated in this study presented highly homologous genotypes, being mostly monomorphic in all loci and most of them sharing the same alleles with very little genetic variability. The observed (Ho) and expected (He) heterozygosity in guanaco and vicuña mites remained within expected parameters, suggesting that they were in HWE. In HWE populations, allele and genotype frequencies are assumed to remain constant from generation to generation in the absence of other evolutionary influences (migration, mutation, selection, gene drift), suggesting that the mange epidemic described here originated from a single source and a single introduction event. Low genetic diversity is common in newly introduced pathogens [24] and consistent with the rapid spread of an emerging pathogen [25]. Unfortunately, at the time of our study there were no llama with mange in the area, which precluded us from further exploring this species as a source of mite introduction. Future studies should apply advanced molecular techniques (e.g. single nucleotide polymorphisms) to clarify the phylogenetic relationships, host preference of mites, mechanisms of propagation, and the source and origin of infestations [6, 56]. Such studies have informed on domestic animal sources in wildlife outbreaks [57] as well as on transmission between domestic and wild animals [58].

Regardless of the origin of the outbreak reported here, the most efficient management approach going forward would be to avoid the presence of livestock within protected areas and to enforce adequate disease prevention and control practices in conservation units that allow livestock grazing. Health risks associated with movements of livestock near national parks are rarely considered in Argentina, and there is little communication between the conservation and agriculture sectors. Thus, livestock incentive programs like the one described here occur under a totally separate set of priorities, agencies, and legislation, with no overlap or consultation with the environmental sector. Moreover, sarcoptic mange is not a mandatory reportable disease in Argentina, so records on species and areas affected are not available.

The establishment of a llama breeding program, which included their introduction to the SGPR without previous consideration of the disease risks due to their taxonomic proximity with the native camelids protected there, plus the discontinuation of veterinary care for introduced animals, carried a high cost for vicuña and guanaco in SGNP. Despite this being a protected area, since the outbreak the local extinction of wild camelids in SGNP is a real possibility, with potential cascading ecological impacts at the community and ecosystem levels. Only science-based, comprehensive and multi-sectorial policies that bridge the environment and livestock sectors can herald a better future for the health of all species.

## Conclusions

Sarcoptic mange had an epidemic behavior with a devastating effect on wild camelids at SGNP. At the end of this study, a scenario of high risk for local extinction of vicuña and guanaco in this protected area was evident. Several factors may have contributed to the rapid spread of mange in SGNP, including a high sensitivity of the animals to the mite evidenced by a severe clinical form of the disease; the social nature and gregarious behavior of camelids; and the initial high densities of camelids in the park, which would have favored contact between individuals and significant spatio-temporal overlap between healthy and sick animals. Mange infection and high susceptibility to puma predation were determining factors in the population collapse observed.

A series of considerations support the hypothesis of the origin of the outbreak in introduced llama: a) from the interviews, two sites of spatio-temporal overlap between introduced llama and wild camelids were detected within and around SGBR; b) there were temporal coincidences between the launch of llama production in San Juan (2009-2014) and the detection of the first cases of mange in native camelids in the park (2014); c) sarcoptic mange is a frequent problem in llama in at least one of the sites of origin of the introduced animals (Cieneguillas); d) mange was diagnosed in some llama entering San Juan from Cieneguillas, and it is possible that further unnoticed cases occurred, either due to lack of reports or subclinical and/or mild infestations; e) interviews suggest that mange has not been a problem in livestock in the last 20 years in SGNP’s area of influence, and no outbreaks of mange have been reported in native camelids in the area, at least in the last 50 years; f) the genetic characteristics of the mites recovered from guanaco and vicuña suggest that it was a recent introduction, with no time to co-evolve with SGNP wild camelids, supporting that the mite is exogenous to the affected population; g) the aggressive and epidemic behavior in SGNP vicuña and guanaco suggests no prior contact with the disease (“naïve” population).

The transmission of diseases between wild and domestic animals will be an increasing challenge at the interface. In Argentina, sarcoptic mange is not notifiable in livestock, but should be considered by the national veterinary service so that efficient disease control mechanisms are implemented in interprovincial animal movements, particularly when they involve protected areas. Proper sanitary management of domestic animals will always be a more reasonable and feasible strategy than trying to contain epidemics in wild populations. With the loss of the largest and main herbivores in the SGNP system, large ecosystem-wide changes are expected in the park. Long-term monitoring will provide valuable information to assess the resilience of the system in response to disease-driven extinction of key species.

## Acknowledgments

We are grateful to personnel of San Guillermo National Park and Administración de Parques Nacionales for logistical support during fieldwork. We extend our gratitude to technicians from the Secretaría de Ambiente y Desarrollo Sustentable of San Juan Province for the information provided, and to llama producers for their collaboration. We thank the Master’s Program in Wildlife Management from Facultad de Ciencas Exactas, Físicas y Naturales de la Universidad de Córdoba Argentina, for assisting with the export of samples. Special thanks to J. Riner and K. Clark for their assistance in molecular analysis, and to veterinarian R. Camera and those interviewed during this study for sharing their knowledge. We thank volunteers F. Vacaflor and P. Muñoz for assistance in the field.

## Supporting Information

**Table S1: Mange detection survey effort (km) per transect in each month. The Agüita del Indio transect was not surveyed on May 2017 due to road blockage by excessive snow.**

**Table S2: Odds ratio (level A relative to level B) of different variables with regards to the occurrence of mange in live vicuña (*Vicugna vicugna*). Asterisks indicate significant differences among levels.**

**Table S3: Odds ratio (level A relative to level B) of different variables with regards to the occurrence of different clinical stages of mange in live vicuña (Vicugna vicugna). Asterisks indicate significant differences among levels.**

**Figure S1: Histology from vicuña (*Vicugna vicugna*) and guanaco (*Lama guanicoe*) carcasses revealed typical sarcoptic mange lesions with abundant mites in all specimens. (A) Epidermis with marked hyperkeratosis: 1- Keratin sheets, 2-Presence of parasites in the stratum corneum, 3- Mixed inflammatory infiltrate in the dermis (10x). (B) 1- Mite in the hyperplastic stratum corneum, 2- dermal collagen sclerosis, 3- sebaceous gland hyperplasia (20x). (C) 1-Plasmacytes, 2- Necrotic material and pustule. Necrotic remains of neutrophils in stratum corneum, 3- Macrophage (40 x). (D) Epidermis with marked hyperkeratosis, 1- Presence of parasite in the stratum corneum, 2- Keratin sheets and remains of scab material, 3- Mixed inflammatory infiltrate in the dermis (10 x).**

**Figure S2: A and B: Llama (*Lama glama*) with alopecic scaling and crusts on forelimbs indicative of sarcoptic mange (Photo: M. Ciallela). These photographs were taken upon arrival of llama to Rodeo (San Juan province) from Cieneguillas (Jujuy province) in 2009.**

**Table S4: Chi-square test summary comparing observed and expected heterozygosis for the Hardy-Weinberg equilibrium in mites from guanaco (Lama guanicoe) and vicuña (Vicugna vicugna).**

